# Can metabarcoding resolve intraspecific genetic diversity changes to environmental stressors? A test case using river macrozoobenthos

**DOI:** 10.1101/2020.03.08.982561

**Authors:** Vera Marie Alida Zizka, Martina Weiss, Florian Leese

## Abstract

Genetic diversity is the most basal level of biodiversity and determines the evolutionary capacity of species to adapt to changing environments, yet it is typically neglected in routine biomonitoring and stressor impact assessment. For a comprehensive analysis of stressor impacts on genetic diversity, it is necessary to assess genetic variants simultaneously in many individuals and species. Such an assessment is not as straight-forward and usually limited to one or few individual species. However, nowadays species diversity can be assessed by analysing thousands of individuals of a community simultaneously with DNA metabarcoding. Recent bioinformatic advances also allow for the extraction of exact sequence variants (ESVs or haplotypes) in addition to Operational Taxonomic Units (OTUs). By using this new capability, we here evaluated if the analysis of mitochondrial genetic diversity in addition to species diversity can provide insights into responses of stream macrozoobenthic communities to environmental stressors. For this purpose, we analysed macroinvertebrate bulk samples of three German river systems with different stressor levels using DNA metabarcoding. While OTU and haplotype number were negatively correlated with stressor impact, this association was not as clear when looking at haplotype diversity. Here, stressor responses were only found for sensitive EPT (Ephemeroptera, Plecoptera, Trichoptera) taxa, and those exceedingly resistant to organic stress. An increase in haplotype number per OTU and haplotype diversity of sensitive taxa was observed with an increase in ecosystem quality and stability, while the opposite pattern was detected for pollution resistant taxa. However, this pattern was less prominent than expected based on the strong differences in stressor intensity between sites. To compare genetic diversity among river systems, only OTUs could be used, which were present in all systems. As OTU composition differed strongly between the rivers, this led to the exclusion of a high number of OTUs, especially in diverse river systems of good quality, which potentially diminished the genetic diversity patterns. To better understand responses of genetic diversity to environmental stressors for example in river ecosystems, it would be important to increase OTU overlap between sites of comparisons, e.g. by sampling a narrower stressor gradient, and to perform calibrated studies controlling for the number and individual genotypes. However, this pioneer study shows that the extraction of haplotypes from DNA metabarcoding datasets is a promising tool to simultaneously assess mitochondrial genetic diversity changes in response to environmental impacts for a metacommunity.

## Introduction

Degradation, pollution, and exploitation of freshwater ecosystems has resulted in a drastic decline of biodiversity (Vörösmarty et al., 2010; WWF, 2018). The magnitude of biodiversity loss depends on stressor intensities as well as on resistance and resilience of the biotic communities (Dobson et al., 2006; Elmqvist et al., 2003; Vörösmarty et al., 2010). So far, degradation and recovery processes have mostly been studied at the level of species diversity (alpha diversity). However, the underlying genetic diversity within species is an essential variable to consider in this context, as it determines the evolutionary capacity of a species to adapt to changing environments. Genetic diversity in ecosystems is influenced by four major processes: mutation, drift, migration, and selection (Vellend and Geber, 2005). A high level of genetic variation is assumed to occur in intact and stable ecosystems, where effective population sizes are large and relatively constant over time. Under stressor impacts, genetic diversity declines (‘genetic erosion hypothesis’) primarily due to reduced population sizes leading to enhanced genetic drift (Amos and Balmford, 2001; Reusch et al., 2005; Reynolds et al., 2012; Ribeiro and Lopes, 2013; van Straalen and Timmermans, 2002). Genetic diversity is the most basal level of biodiversity and typically the first to decrease under, and the last to regenerate after stressor impact. It consequently provides a proxy for ecological processes long before, or even if never visible on the species diversity level (Bazin et al., 2006; Guttman, 1994; Hughes et al., 2008; Reynolds et al., 2012; Vellend and Geber, 2005). In conservation biology, genetic diversity is therefore used as a measure for population and habitat stability. This information is typically neglected in the legally binding species assessment, or species diversity of a habitat and is regarded as a proxy for genetic diversity (Vellend, 2005; Vellend and Geber, 2005). However, this concept has been rarely tested and has been questioned for example by Taberlet et al. (2012) who showed that for alpine plant species genetic diversity is not correlated with alpha diversity.

Nowadays, alpha diversity can be assessed with great resolution using DNA metabarcoding (e.g. Deiner et al., 2016; Hänfling et al., 2016; Macher et al., 2018). With these data, stressor impacts can be analysed simultaneously for many taxa not distinguishable by morphological determination methods (Bagley et al., 2019; Beermann et al., 2018; Pfrender et al., 2010; Theissinger et al., 2019). Typically, DNA metabarcoding infers responses at Operational Taxonomic Unit (OTU) level. In most cases, distance-based thresholds are used to define OTUs with the aim that these reflect as closely as possible biological species. While OTU classification can drastically improve the taxonomic and ecological resolution in comparison to classical morphological taxa assignment (Beermann et al., 2018; Macher et al., 2016; Sturmbauer et al., 1999), the level of intraspecific genetic diversity still goes unnoticed, when distance-based clustering is used to analyse the sequencing data. As an alternative, bioinformatic denoising approaches can be used to obtain ‘exact sequence variants’ (ESVs) (Callahan et al., 2016; Frøslev et al., 2017). Here, additional bioinformatic filtering steps are used, that allow to separate biological template sequences from noisy reads caused by PCR and sequencing errors. With ESVs it is possible to explore intraspecific genetic diversity patterns in eukaryotes (Elbrecht et al., 2018; Tsuji et al., 2019; Turon et al., 2019). For animals, commonly a 658-bp fragment of the mitochondrial cytochrome c oxidase I gene (COI) is used for DNA barcoding and a shorter part of this for metabarcoding (Hebert et al., 2003). While the use of haploid, maternally-inherited mitochondrial markers has limitations for detailed population genetic analyses (Ballard and Whitlock, 2004; Leese and Held, 2011), its utility to infer insights into geographic structure and population diversity, and thereby ecological processes acting at local or regional level, has often been demonstrated (e.g. Pauls et al., 2006; Weiss and Leese, 2016; Witt and Hebert, 2000). Furthermore, extracting intraspecific haplotype variation data from COI metabarcoding datasets that typically operate only on species diversity level is regarded as a promising tool to gain better understanding of metacommunity structure and stressor impacts, to eventually manage natural communities more efficiently than possible with species diversity data alone (Adams et al., 2019; Geist, 2011; Hughes et al., 2008; Reusch and Hughes, 2006; Reynolds et al., 2012).

In our study, we want to further evaluate the potential of ESV, i.e. COI haplotype data in addition to OTU data obtained from environmental bulk sample metabarcoding datasets, for analysing the impact of stressors on macrozoobenthos (MZB) communities in river ecosystems. MZB organisms play a key role in freshwater ecosystem functionality and include a wide range of taxonomic groups with often narrow and specific demands with respect to habitat conditions (Jackson and Füreder, 2006; Usseglio-Polatera et al., 2000; Wallace and Webster, 1996). As river ecosystems, three German rivers systems with differing stressor impacts were chosen: Emscher – high stress, Ennepe – moderate stress, and Sieg – low stress. The main branch of the Emscher is an urban stream in the Ruhr Metropolis and has been used as an open sewage channel for the past hundred years. It is now part of one of the biggest restoration projects in Europe, yet impacts on stream biota are pervasive in most parts. Sample sites in this stream were chosen to be in conditions with variable stressor inflow, ranging from completely restored sites, to canalized sites with purification plant inflow, and to sites in unrestored sewage channels. Heterogeneous conditions and stressor impacts are also present in the Ennepe, and include near-natural rural sites, urban sites stressed through occasional stormwater retention basin overflow, and sites with sewage treatment plant inflow. In comparison, the river Sieg is considered as a stable, near-natural river system with a good ecological and chemical status. Punctual stressor inflow is present through rainwater retention basins but not immediately at sampling sites. By comparing communities between the different streams, we want to test, if species and genetic diversity is correlated with present stressor gradients. Following predictions from the ecological habitat concept, which links presence and abundance of species over time to the available resources (see e.g. Van Dyck, 2012), we expect OTU diversity to be highest in the river Sieg, moderate in river Ennepe, and lowest in river Emscher. According to the genetic erosion hypothesis, we predict haplotype diversity to covary with OTU richness. We expect highest haplotype number and diversity at the river Sieg due to its long-time stable good ecological conditions supporting large and stable population sizes. Lower values are expected at the river Ennepe, where communities are regularly affected by organic stressor influx and thus recurrent population decline, resulting in higher genetic drift. The lowest values are expected for river Emscher due to the complete erasure of MZB diversity in the history of this river system caused by the usage as sewage transport system, and the still prevalent stress level in many parts. We predict different patterns for taxa sensitive (EPT – Ephemeroptera, Plecoptera, Trichoptera) or resistant to organic pollution (PR – ‘Pollution Resistant’ – Arhynchobdellida, Enchytraeida, Haplotaxida, Isopoda, Rhynchobdellida). Specifically, we assume that OTU and haplotype diversity for EPT taxa will decline with increasing stress because of declining population sizes, and eventual local species extinction. In contrast we expect that PR taxa will show opposing patterns, because of their resistance to organic pollution and their ability to rather use organic pollutants as a resource, potentially facilitating large population sizes (Friberg et al., 2010; Ribeiro and Lopes, 2013; Smith et al., 2007).

## Material and Methods

### Sampling

Macroinvertebrates were sampled according to Water Framework Directive-compliant protocols (Meier et al. 2006) at six sites in the rivers Emscher and Sieg in autumn 2016 & 2017, and spring 2017 & 2018 (Figure 1). In short, kick-net sampling of different habitats with 20 subsamples in the Sieg, and ten subsamples in the Emscher due to fewer available microhabitats, was executed. The seven sites at the Ennepe were sampled in autumn 2017 and spring 2017 & 2018 in a similar fashion with ten subsamples. Subsamples were pooled, large parts of substrate discarded (e.g. stones, leaves, small branches), and samples, including macrozoobenthic specimens and remaining substrate, were transferred to 1L bottles filled up with ethanol. Approximately 1/3 of bottle volume was filled with the sample and 2/3 with 96% technical ethanol. If volume of sampled material was too large, it was divided into multiple bottles. Samples were transported to the laboratory, and old ethanol was replaced with new 96 % technical ethanol on the same day.

**Figure 1:**
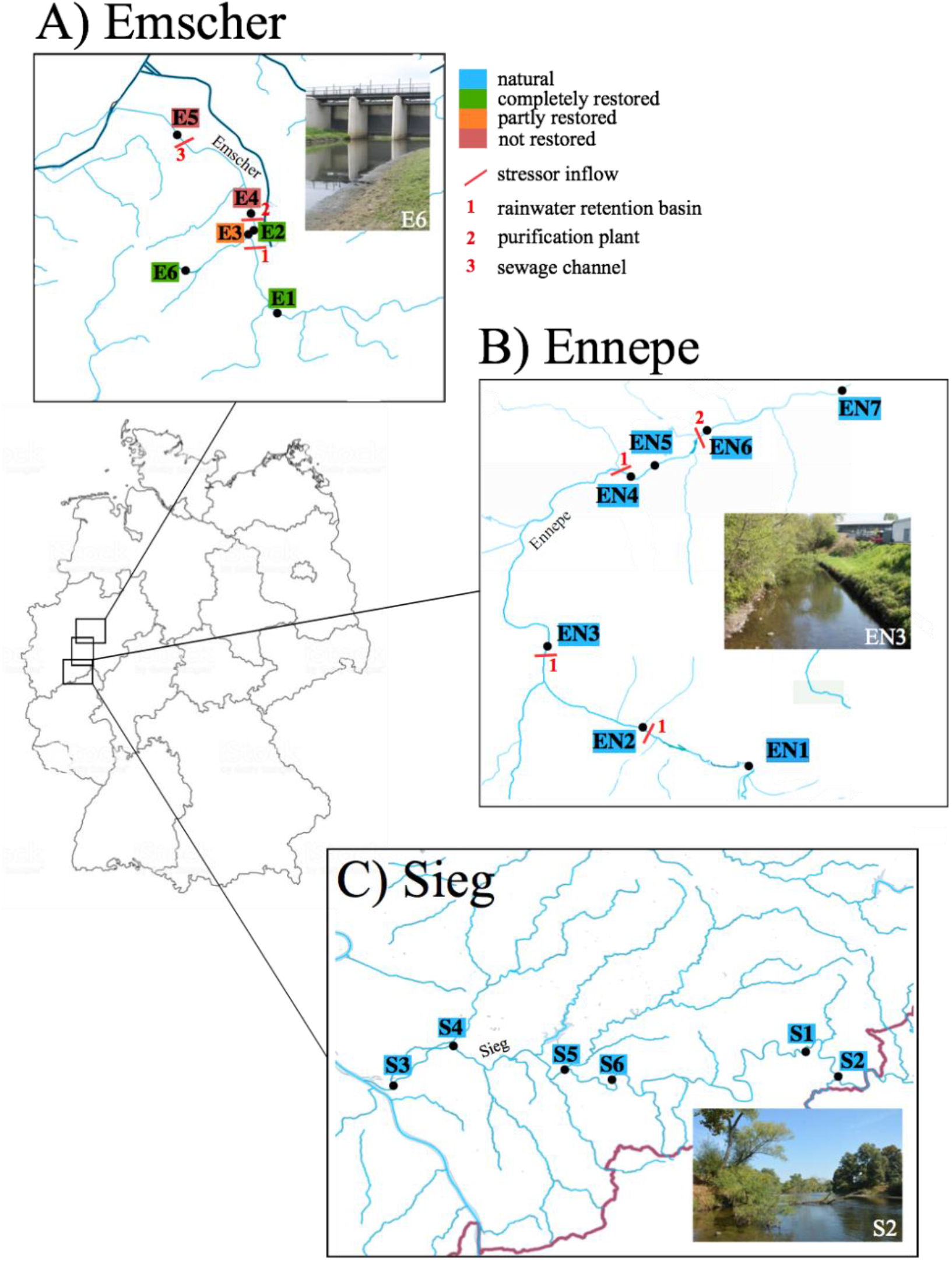
Sample sites at river A) Emscher, B) Ennepe and C) Sieg.

### Laboratory Protocols

Samples were examined under a binocular (Leica S6E) to separate individuals from substrate. Substrate was discarded and individuals were counted and separated in two size categories (size class A: ≤ 25 mm, size class B: ≥ 25 mm) (see Elbrecht et al. 2017b for the procedure). Individuals of the two size classes were dried in petri dishes overnight and homogenized to fine powder with an IKA Ultra Turrax Tube Disperser (BMT-20-S-M sterile tubes, full speed for 30 min). Two times half of a spatula (~20 mg) was transferred into two Eppendorf tubes and 600 μl TNES buffer and 15 μl Proteinase K (10 mg/ml) were added per tube and incubated over night at 36 °C shaking at 250 rpm in a Thermoshaker (ThermoMixer C, Eppendorf). A salt extraction protocol (after Sunnucks and Hales 1996, adjusted as in Weiss and Leese, 2016) was used to isolate DNA from powder. After DNA extraction and subsequent RNA digestion (1 μl RNase A (ThermoFisher Scientific) per sample, incubated at 37 °C for 30 min), samples were cleaned up (NucleoSpin gel and PCR clean up kit, Macherey-Nagel), and size groups per sample were pooled according to specimen numbers per group. DNA was quantified with a Qubit Fluorometer (ds DNA BR Assay kit, Thermo Fisher Scientific, Beverly, USA) and adjusted to 25 ng/μl. A two-step PCR (for futher information see Zizka et al., 2019) was conducted with one technical (PCR) replicate per sample. The universal BF2/BR2 primers (Elbrecht and Leese, 2017) were used. PCR reactions included 1× PCR buffer (including 2.5 mM Mg_2+_), 0.2 mM dNTPs, 0.5 μM of each primer, 0.025 U/L of HotMaster Taq (5 Prime, Gaithersburg, MD, USA), and 1 μL DNA template filled up with HPLC H_2_O to a total volume of 50 μL. PCR conditions were: 94 °C for 180 s; 25 cycles of 94 °C for 30 s, 50 °C for 30 s, and 65 °C for 150 s; followed by a final elongation of 65 °C for 5 min in a Thermocycler (Biometra TAdvanced). A left-sided size selection was conducted per sample with magnetic SpriSelect beads (Beckman Coulter, Krefeld, Germany), using a ratio of 0.76× to remove small fragments (primers, primer dimers). DNA concentration after PCR was measured on a Fragment Analyzer (Advanced Analytical, Ankery, USA) and samples were pooled equimolarly. Library pools were sent for paired-end sequencing to Eurofins (Constance, Germany) on four Illumina MiSeq runs (2×250 bp paired-end v2 kit), one for each sampling season.

### Data Analysis

Sequences were analysed with JAMP-0.67 (https://github.com/VascoElbrecht/JAMP) including demultiplexing of data, paired-end-merging, and primer trimming following standard settings. For haplotype extraction only reads of expected fragment length (421 bp) were included. A strict quality filtering was applied (maximal expected error *max_ee = 0.3*) and reads with an abundance < 0.003 % in a sample were excluded from the dataset. The algorithm Unoise3 (Edgar, 2016) implemented in JAMP, was used to denoise the dataset (*alpha = 5*) and to separate common haplotypes from chimeras and sequencing noise. The denoising approach is based on the assumption, that high abundant unique reads (centroids) are real sequences amplified from the biological template. Defined by distance (d), other unique sequences (neighbours) are grouped around these high abundant sequences. Based on the Levenshtein distance and abundance (defined by *alpha*), neighbours showing a small difference and abundance compared to the centroid are predicted to be erroneous. Unoise3 has been shown to be particularly efficient in past studies under controlled conditions (Tsuij et al. 2020). Denoised reads were assigned to OTUs (clustered by 3 % distance) and the number of sequences per haplotype were determined in each sample. As a further filtering step, OTUs with an abundance below 0.01 % (*OTUmin = 0.01*) in at least one sample and haplotypes with an abundance below 0.003 % (*minhaplosize = 0.003*) in at least one sample were discarded (see Elbrecht et al. 2018 for detailed explanation). This step was included, to filter also low abundant unique sequences, which are not integrated in the filtering through *alpha*. Taxonomic assignment of haplotypes was conducted through a comparison with the database BOLD (Ratnasingham and Hebert, 2007) using an in-house python script (available on request). Haplotypes with similarity < 95 % to a deposited sequence in the database were excluded from further analysis to prevent incorrect assignments potentially leading to the assessment of erroneous diversity patterns. Read numbers per haplotype of technical PCR replicates were fused and the average was calculated. Further analyses were carried out with the average read number per haplotype. To assess haplotype richness per OTU, we used count data. However, in order to approximate also traditional population genetic measures, we calculated haplotype and nucleotide diversity per sample site and season with Arlequin 3.5 (Excoffier and Lischer, 2010) using read depths as a proxy for haplotype abundance. Data were not normally distributed and therefore the non-parametric Kruskal-Wallis test was used to check for effects of river system on diversity variables. A post-hoc Dunn test (package dunn.test()), Dinno, 2017) was used to conduct pairwise comparisons for significant differences. All statistical analysis were conducted in R (R Development Core Team, 2008). Table modification and figure preparation was done using the packages vegan (Oksanen et al., 2019), tidyverse (Wickham et al., 2019) and ggplot2 (Wickham, 2016) implemented in R.

## Results

After denoising and abundance filtering, on average 29,063 reads were present in Emscher samples, 36,289 reads in Sieg samples and 51,981 reads in Ennepe samples (Table S1). Because filtering thresholds were based on relative abundances (see Material and Methods, Data Analysis), reads per sample were not adjusted to uniform numbers. Samples contained 228 – 694 haplotypes, which clustered into 70 – 155 OTUs. OTU and haplotype number was higher at river Ennepe and Sieg than at river Emscher (p < 0.01, Fig. 2). A high number of unique haplotypes per sample site was detected in all river systems (Fig. S7). Splitting the dataset in EPT (Ephemeroptera, Plecoptera, Trichoptera) and PR (‘Pollution Resistant’) taxa revealed 13 - 273 haplotypes clustered into 2 – 46 OTUs (zeros excluded, e.g. E5) for EPT taxa, and 8 – 257 haplotypes clustered into 4 - 55 OTUs for PR taxa. As no plecopterans were found at river Emscher, only ET taxa could be analysed for this river. Total OTU and haplotype number of EPT taxa was affected by the river system with more OTUs and haplotypes at Ennepe and Sieg than at Emscher (p < 0.001). Sample sites E4 and E5 showed remarkably high OTU and haplotype numbers assigned to PR taxa compared to all other sample sites. However, no effect of the river system was detected on PR taxa (haplotype number p = 0.09) (Fig. 2). Number of counted individuals before laboratory processing differed between all river systems (p < 0.001) and seasons (p < 0.05) (Table S2). Within streams, individual numbers did not significantly differ between sampling sites. However, no correlation was detected between total specimen number per site and season, and average haplotype number per OTU (Fig. S1).

**Fig. 2:**
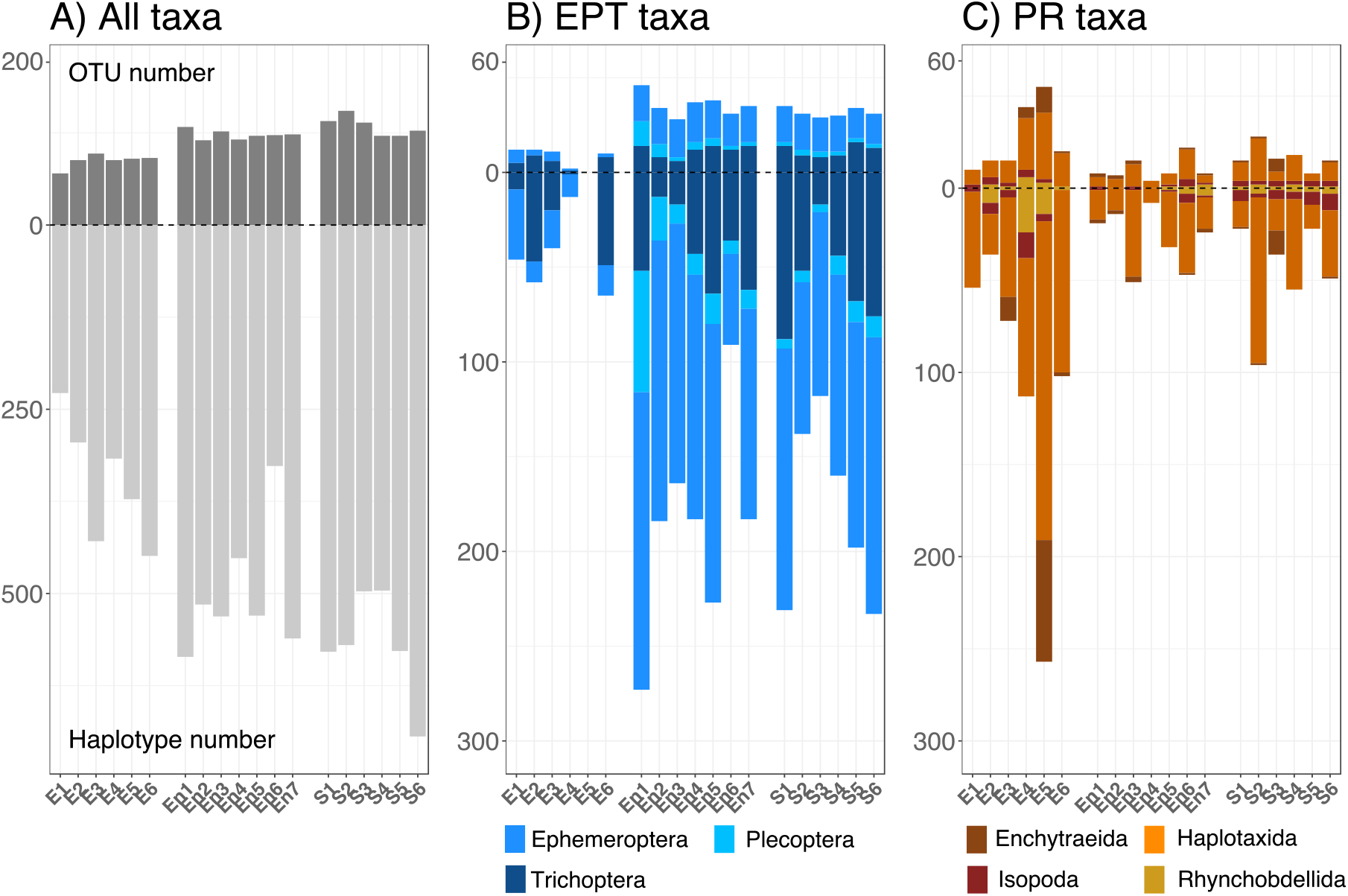
Total OTU (upper part) and haplotype number (lower part) summed across all seasons for all river systems (Em 1-6 – Emscher, En 1-7 – Ennepe 1-6, S - Sieg). A) All benthic macroinvertebrate taxa assigned to a reference in BOLD with > 95 %; B) EPT taxa (Ephemeroptera, Plecoptera, Trichoptera); C) PR taxa (Pollution Resistant → Enchytraeida, Haplotaxida, Isopoda, Rhynchobdellida).

To compare average haplotype number per OTU and haplotype diversity between river systems, we searched for OTUs present in most samples. The five most common OTUs, all occurring at more than 50 % of the analysed samples, were: OTU 2 (*Baetis rhodani*, 63 %), OTU 9 (*Asellus aquaticus*, 52 %), OTU 12 (*Stylodrilus heringianus*, 63 %), OTU 65 (*Esolus parallelepipedus*, 55 %), and OTU 107 (*Microtendipes pedellus*, 52 %). To further increase the number of shared OTUs between sites, samples collected at different seasons were merged (E1-E6, En1-En7, S1-S6). By this, we identified six OTUs occurring in more than 80 % of all sites (OTU 2: *Baetis rhodani*, 84 %; OTU 5: *Orthocladius*, 89 %, OTU 9: *Asellus aquaticus*, 84 %; OTU 45: *Orthocladius* sp., 84 %; OTU 67: *Tanytarsus eminulus*, 84 %, OTU 107: *Microtendipes pedellus*, 95 %). To increase the number of OTUs for analyses, all OTUs present in at least one of the samples per river system were included, resulting in four different data sets (Em-En-S: 78 shared OTUs, Em-En: 110 shared OTUs, Em-S: 125 shared OTUs, En-S: 155 shared OTUs). Per dataset > 47 % of shared OTUs were assigned to dipterans, of which the majority (> 90 %) were chironomids (Figure S2). Comparisons of average haplotype number per OTU and haplotype diversity revealed no differences between river systems for all four datasets when all taxa were included (Fig. S3). Dividing the datasets into OTUs assigned to EPT (pollution sensitive) and PR (pollution resistant) taxa revealed a significant effect of the river system on the average haplotype number per shared OTU when comparing all three river systems (EPT: p < 0.05, PR: p < 0.05) (Fig. 3). Ennepe and Sieg showed a higher average haplotype number per OTU for EPT taxa than the Emscher (En: 4; S: 3.3; Em: 2.7, p < 0.05), while the ratio for PR taxa was higher at Emscher (3.1) than at the other two rivers (En: 2.7, S: 2.6). When comparing only shared PR OTUs between Emscher and Sieg, more haplotypes per OTU were found at the Emscher (Em: 3.2; S: 2.4, p < 0.05). Observations on PR taxa also showed a higher haplotype diversity at the Emscher in comparison to both other streams (Em: 0.374, En: 0.303, S: 0.249). When comparing shared OTUs only between Emscher and Sieg, a significantly higher haplotype diversity of PR taxa was observed at the Emscher (Em: 0.3338; S: 0.1934, p < 0.05) (Fig. 3). Detailed information on average haplotype number per OTU and haplotype diversity per sample site are illustrated in figure S3 (average haplotype number per OTU) and S4 (nucleotide diversity).

**Fig. 3:**
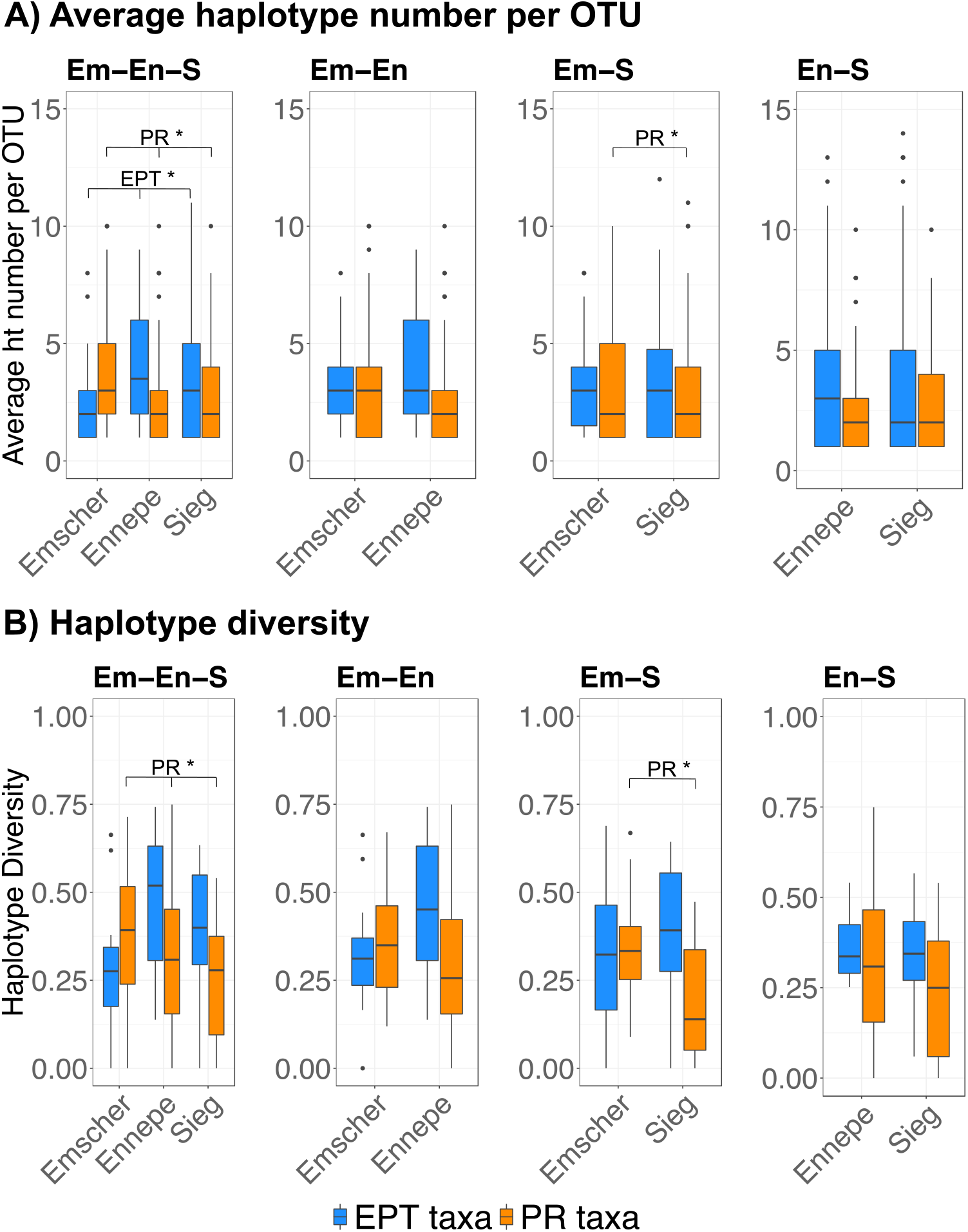
A) Average haplotype number per OTU for the four datasets of shared OTUs. B) Haplotype diversity for the four datasets of shared OTUs. Sensitive (EPT) and pollution resistant (‘PR’) taxa are shown. * indicates significant difference between regarded groups.

Further, the comparison of shared OTUs between the three river systems revealed an effect of river system on nucleotide diversity for EPT taxa (p < 0.05). Average nucleotide diversity of taxa was higher at river Ennepe (0.00215) and Sieg (0.00266) than at the Emscher (0.00104). In contrast, PR taxa showed a higher nucleotide diversity at the Emscher (0.00143) than at the Ennepe (0.00215), but only when comparing OTUs shared between those two rivers (p < 0.05, Fig. S6).

We plotted total OTU number assigned to EPT and PR taxa against the average haplotype number per OTU and sample site for the four datasets of shared OTUs (Fig. 4 A-H) to test if OTU and genetic diversity are linked and indirectly, if ecosystem quality effects genetic variability. A significant correlation was observed for ET taxa comparing all three river systems with a clear increase from Emscher to Sieg and Ennepe (Fig. 4 A). A weaker correlation was observed for EPT taxa comparing Ennepe and Sieg (Fig. 4 D). In addition, correlations were significant for PR taxa comparing Emscher, Ennepe and Sieg (Fig. 4 G) as well as Emscher and Sieg (Fig. 4 G) which were mainly driven by a few samples of extremely high OTU number. A clear separation of river systems according to total number of ET taxa (x axis) was visible comparing all three river systems (Fig. 4 A) and in comparisons between Emscher and Ennepe (Fig. 4 B) and Emscher and Sieg (Fig. 4 C). The separation was most distinct between Emscher and Ennepe. Separation of river systems due to total number of PR taxa (x axis Fig. E-H) was less distinct than separation based on total ET taxa.

**Fig. 4:**
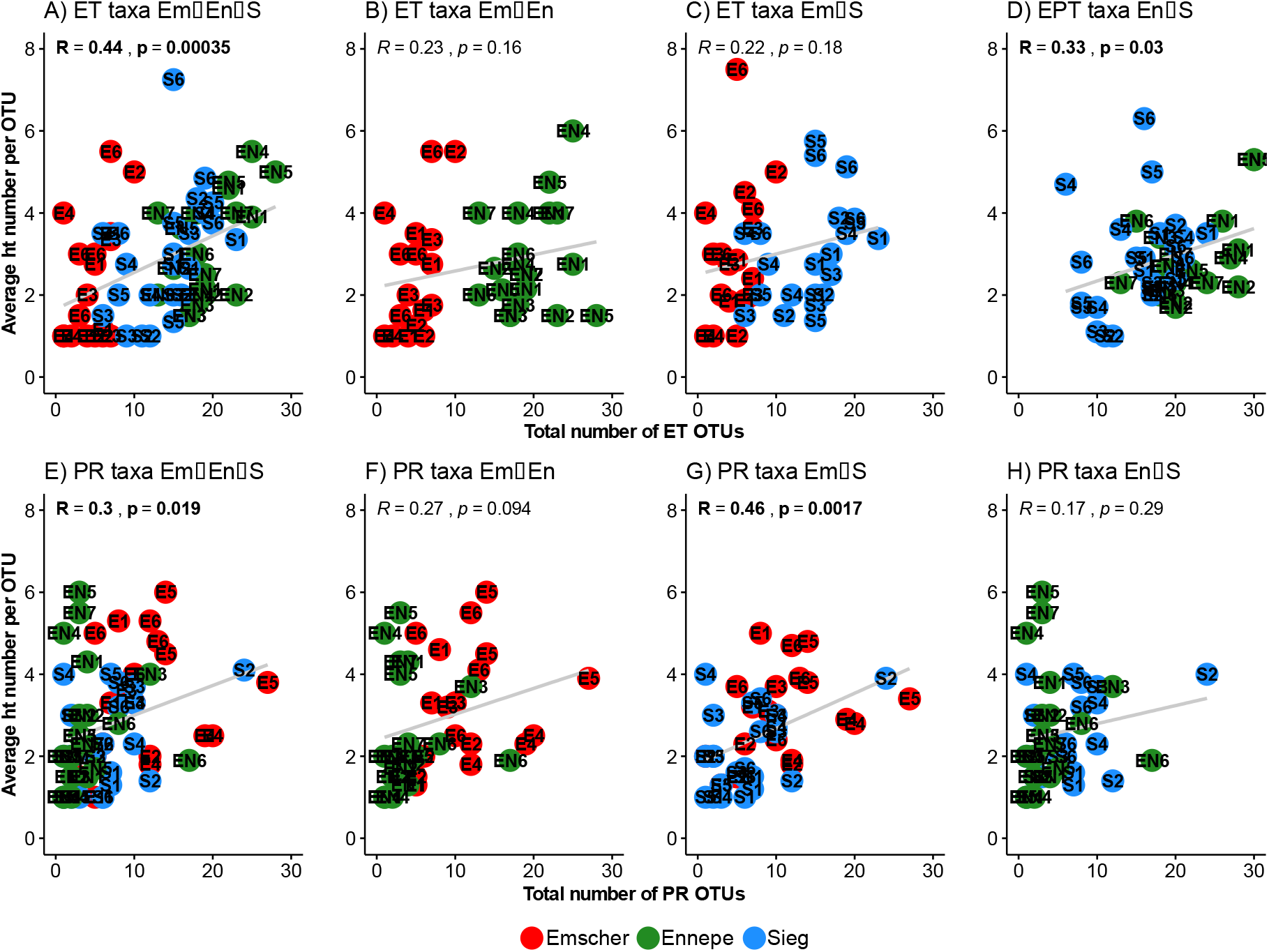
Correlation analysis of total number of OTUs assigned to EPT/PR taxa and average haplotype number per OTU for those groups. A-D four datasets of shared OTUs for EPT taxa. E-H four datasets of shared OTUs for PR taxa.

## Discussion

We expected a general effect of stressors on alpha (OTU) diversity and intraspecific genetic diversity in the three river systems. In line with these expectations, the heavily impacted river Emscher showed the lowest OTU number compared with the other two systems. However, no significant differences between Ennepe and Sieg were detected, and an even higher number of highly pollution-sensitive stoneflies (Plecoptera) was found at the stronger impacted river Ennepe. Stressor effects at this river therefore seem to be less intense than expected with no obvious effects on MZB communities. As expected, haplotype numbers per sample site were lower in river Emscher than in Emscher and Sieg, especially for pollution-sensitive mayfly and caddisfly taxa. Vice versa, pollution resistant (‘PR’) taxa had higher haplotype diversity in the river Emscher. Genetic diversity estimates inferred via average haplotype number per OTU, haplotype diversity as well as nucleotide diversity revealed higher values at river Sieg and Ennepe compared the Emscher but no differences between the former two rivers and therefore supported results based on OTU and haplotype number. In this case, the hypothesis that sustainable good ecological conditions and stability at river Sieg induced stable and large population size, favouring a high level of genetic diversity in sensitive MZB communities was supported (Reynolds et al., 2012; Ribeiro and Lopes, 2013). Results further indicate similar conditions at the Ennepe with lower stressor intensity than expected and no distinct effect on the sensitive community composition of MZB taxa and according genetic variability. The Em-En-S dataset further underlines a strong correlation between average haplotype number per OTU and total number of ET taxa, further emphasizing a linkage between species (OTU) diversity and genetic variability (Vellend and Geber, 2005). Assuming a higher number of sensitive ET taxa in ecologically intact streams, the correlation also links local habitat condition with genetic diversity, increasing from Emscher to Sieg and Ennepe. For the other datasets (Fig. S2) no significant differences in genetic diversity could be observed for E(P)T taxa. Beside actual biological signal, this is most likely due to the insufficiency of underlying datasets or methodological problems that will be discussed in the following paragraphs (from paragraph ‘ *OTU overlap’* on). Pollution resistant taxa showed a higher average haplotype number per OTU and haplotype diversity at the Emscher than the other two systems when all three rivers were compared. This supports initial assumptions about low competition pressure and increased population growth of those taxa in stressed systems (Friberg et al., 2010; Gaufin and Tarzwell, 1952; Smith et al., 2007). Detected patterns comparing all the river systems are also supported by significant differences in genetic diversity between Emscher and Sieg and correlations between total number of PR taxa and genetic diversity.

The data presented hold great potential to inform on metacommunity and metapopulation structure and processes. However, several limitations need to be considered when further testing and applying the approach:

### OTU overlap

The fact that no significant differences between average haplotype number per OTU and haplotype diversity were found for the other comparisons of E(P)T taxa (pairwise comparisons of all rivers) is most probably affected by the small overlap of OTUs shared between rivers. The highly heterogeneous, site-specific stressor levels and variable time since restoration at Emscher further inflated variation in OTU composition, thereby limiting statistical power for comparisons of genetic variation based on highly frequent OTUs. Datasets had to be reduced to OTUs at least present in one sample site of compared river systems to circumvent differences in genetic variability due to species specific variations. However, this approach also excludes taxa unique to specific sites, limiting analysis of metacommunities at highly diverse stream ecosystems (Sieg, Ennepe), and also limiting number of metapopulations to study with respect to mitochondrial genetic diversity.

### Mitochondrial single-gene marker

Beside the limitations due to a small overlap in OTUs between river systems, the utility of the mitochondrial cytochrome c Oxidase I (COI) marker as a measure of intraspecific genetic diversity and variability could have deflated signal strength. Even if various studies have successfully applied this marker for even more specific population analyses, the use of only a single gene can be misleading – and mitochondrial genes are especially unique due to their haploid structure, the lack of recombination and purely maternal inheritance (Ballard and Whitlock, 2004; Leese and Held, 2011).

### Bioinformatic haplotype extraction

Furthermore, as outlined in previous studies (Elbrecht et al., 2018; Tsuji et al., 2019; Turon et al., 2019), the main challenge in extracting haplotypes from metabarcoding datasets is the separation of ‘real’ environmental sequences from those produced by PCR or sequencing errors. New programs enable the denoising of datasets with the optimisation of filtering steps, to efficiently separate real sequences from erroneous ones. However, a decision has to be made concerning the strictness of filtering. By using high filtering thresholds, erroneous sequences are excluded with a higher probability, but at the same time it is more likely to exclude real sequences of low abundance. In comparison, a lower filtering increases the number of rare real sequences, but also includes a higher number of erroneous sequences into diversity analysis. For the present study, we followed denoising as recommended in Elbrecht et al., 2018, where the genetic variability of benthic macroinvertebrates in Finland streams was investigated, implementing an α-value of 5, which was also applied in Turon et al., 2019. The additional percentual abundance threshold filtering for OTUs and haplotypes was set after suggestions in Elbrecht et al., 2018 and applied in Laini et al., in review. We found a high number of unique haplotypes per sample site (exemplary networks of the two most frequently found EPT and PR taxa shown in Fig. S7), which is similar to other metabarcoding studies (Elbrecht et al., 2018; Laini et al., in review), but exceeds those found in studies based on single specimen barcoding (Lucentini et al., 2011; Weiss and Leese, 2016; Williams et al., 2006). Higher numbers of unique haplotypes could be induced into the dataset through sequencing errors, which would emphasize the application of an even higher filtering threshold on metabarcoding datasets. However, the increased number of haplotypes found through metabarcoding can also be real haplotype variants in one specimen, which cannot be determined through single specimen barcoding due to the underlying sequencing method (Phillips et al., 2019).

### Individual sample size

Lastly, while the strength of the approach is that thousands of specimens were processed at once, there is little control about the individual number of specimens per species or OTU. Comparisons of genetic diversity rely on the number of specimens sampled and future studies should carefully control for this in order to test the reliability and robustness of the approach.

## Conclusion

Our study shows that metabarcoding-derived haplotype information can be utilized to assess effects of environmental variables on intraspecific genetic diversity using the example of three differently-impacted German river systems. Even if the choice of data filtering thresholds is still a trade-off between rare ‘real’ sequences and erroneous, artificially generated ones, we were able to link stressor effects to genetic variability of aquatic macroinvertebrate communities. Sites with good and stable ecological conditions showed higher genetic diversity than stressed sites, which is coupled with OTU diversity. However, due to a low OTU overlap between river systems, genetic diversity analyses were based only on subsets, including all shared OTUs. This subsampling induced the exclusion of variability and ecological specialists, especially at highly diverse sample sites and might skew actual differences. Further analysis of stressor level or restoration events on genetic diversity should include several replicates of similar conditions, presuppose a threshold of overlapping OTUs in compared river systems and involve the control of individuals per OTU. Our study adds to the growing number of studies that highlight the potential to extract haplotype information from metabarcoding datasets for more holistic biodiversity assessments.

## Acknowledgements

This study was supported by the German Federal Ministry of Education and Research project “German Barcode of Life II” (GBOLII) to FL (FKZ 01LI1101 and 01LI1501). We want to thank the Emschergenossenschaft/Lippeverband and Ennepe-Ruhr-Kreis for support during this study and Bianca Peinert for her help in the field and the laboratory. FL and VMAZ are supported by the COST Action CA15219 DNAqua-Net.

## References

Adams, C.I., Knapp, M., Gemmell, N.J., Jeunen, G.-J., Bunce, M., Lamare, M.D., Taylor, H.R., 2019. Beyond Biodiversity: Can Environmental DNA (eDNA) Cut it as a Population Genetics Tool? Genes 10, 192.

Amos, W., Balmford, A., 2001. When does conservation genetics matter? Heredity 87, 257–265. https://doi.org/10.1046/j.1365-2540.2001.00940.x

Bagley, M., Pilgrim, E., Knapp, M., Yoder, C., Santo Domingo, J., Banerji, A., 2019. High-throughput environmental DNA analysis informs a biological assessment of an urban stream. Ecological Indicators 104, 378–389. https://doi.org/10.1016/j.ecolind.2019.04.088

Ballard, J.W.O., Whitlock, M.C., 2004. The incomplete natural history of mitochondria. Molecular Ecology 13, 729–744. https://doi.org/10.1046/j.1365-294X.2003.02063.x

Bazin, E., Glémin, S., Galtier, N., 2006. Population Size Does Not Influence Mitochondrial Genetic Diversity in Animals. Science 312, 570. https://doi.org/10.1126/science.1122033

Beentjes, K.K., Speksnijder, A.G.C.L., Schilthuizen, M., Schaub, B.E.M., van der Hoorn, B.B., 2018. The influence of macroinvertebrate abundance on the assessment of freshwater quality in The Netherlands. MBMG 2, e26744. https://doi.org/10.3897/mbmg.2.26744

Beermann, A.J., Zizka, V.M.A., Elbrecht, V., Baranov, V., Leese, F., 2018. DNA metabarcoding reveals the complex and hidden responses of chironomids to multiple stressors. Environmental Sciences Europe 30, 26. https://doi.org/10.1186/s12302-018-0157-x

Callahan, B.J., McMurdie, P.J., Rosen, M.J., Han, A.W., Johnson, A.J.A., Holmes, S.P., 2016. DADA2: High-resolution sample inference from Illumina amplicon data. Nature Methods 13, 581–583. https://doi.org/10.1038/nmeth.3869

Deiner, K., Fronhofer, E.A., Mächler, E., Walser, J.-C., Altermatt, F., 2016. Environmental DNA reveals that rivers are conveyer belts of biodiversity information. Nature Communications 7, 12544. https://doi.org/10.1038/ncomms12544

Dinno, A., 2017. Package “dunn.test.”

Dobson, A., Lodge, D., Alder, J., Cumming, G.S., Keymer, J., McGlade, J., Mooney, H., Rusak, J.A., Sala, O., Wolters, V., Wall, D., Winfree, R., Xenopoulos, M.A., 2006. Habitat Loss, Trophic Collapse, And The Decline Of Ecosystem Services. Ecology 87, 1915–1924. https://doi.org/10.1890/0012-9658(2006)87[1915:HLTCAT]2.0.CO;2

Edgar, R.C., 2016. UNOISE2: improved error-correction for Illumina 16S and ITS amplicon sequencing. bioRxiv 081257.

Elbrecht, V., Leese, F., 2017. Validation and Development of COI Metabarcoding Primers for Freshwater Macroinvertebrate Bioassessment. Frontiers in Environmental Science 5, 11. https://doi.org/10.3389/fenvs.2017.00011

Elbrecht, V., Vamos, E.E., Steinke, D., Leese, F., 2018. Estimating intraspecific genetic diversity from community DNA metabarcoding data. PeerJ 6, e4644. https://doi.org/10.7717/peerj.4644

Elmqvist, T., Folke, C., Nyström, M., Peterson, G., Bengtsson, J., Walker, B., Norberg, J., 2003. Response diversity, ecosystem change, and resilience. Frontiers in Ecology and the Environment 1, 488–494. https://doi.org/10.1890/1540-9295(2003)001[0488:RDECAR]2.0.CO;2

Excoffier, L., Lischer, H.E.L., 2010. Arlequin suite ver 3.5: a new series of programs to perform population genetics analyses under Linux and Windows. Molecular Ecology Resources 10, 564–567. https://doi.org/10.1111/j.1755-0998.2010.02847.x

Friberg, N., Skriver, J., Larsen, S.E., Pedersen, M.L., Buffagini, A., 2010. Stream macroinvertebrate occurrence along gradients in organic pollution and eutrophication. Freshwater Biology 55, 1405–1419. https://doi.org/10.1111/j.1365-2427.2008.02164.x

Frøslev, T.G., Kjøller, R., Bruun, H.H., Ejrnæs, R., Brunbjerg, A.K., Pietroni, C., Hansen, A.J., 2017. Algorithm for post-clustering curation of DNA amplicon data yields reliable biodiversity estimates. Nature Communications 8, 1188. https://doi.org/10.1038/s41467-017-01312-x

Gaufin, A.R., Tarzwell, C.M., 1952. Aquatic invertebrates as indicators of stream pollution. Public Health Rep 67, 57–64.

Geist, J., 2011. Integrative freshwater ecology and biodiversity conservation. Ecological Indicators 11, 1507–1516. https://doi.org/10.1016/j.ecolind.2011.04.002

Guttman, S.I., 1994. Population genetic structure and ecotoxicology. Environmental Health Perspectives 102, 97–100. https://doi.org/10.1289/ehp.94102s1297

Hänfling, B., Lawson Handley, L., Read, D.S., Hahn, C., Li, J., Nichols, P., Blackman, R.C., Oliver, A., Winfield, I.J., 2016. Environmental DNA metabarcoding of lake fish communities reflects long-term data from established survey methods. Molecular Ecology 25, 3101–3119. https://doi.org/10.1111/mec.13660

Hebert, Paul.D.N., Cywinska, A., Ball, S.L., deWaard, J.R., 2003. Biological identifications through DNA barcodes. Proceedings of the Royal Society of London. Series B: Biological Sciences 270, 313–321. https://doi.org/10.1098/rspb.2002.2218

Hughes, A.R., Inouye, B.D., Johnson, M.T.J., Underwood, N., Vellend, M., 2008. Ecological consequences of genetic diversity. Ecology Letters 11, 609–623. https://doi.org/10.1111/j.1461-0248.2008.01179.x

Jackson, J.K., Füreder, L., 2006. Long-term studies of freshwater macroinvertebrates: a review of the frequency, duration and ecological significance. Freshwater Biology 51, 591–603. https://doi.org/10.1111/j.1365-2427.2006.01503.x

Laini, A., Beermann, A.J., Bolpagni, R., Burgazzi, G., Elbrecht, V., Zizka, V.M.A., Leese, F., Viaroli, P., in review. Metabarcoding improves the detection of nestedness-turnover components of beta diversity in intermitted streams.

Leese, F., Held, C., 2011. Analysing intraspecific genetic variation: a practical guide using mitochondrial DNA and microsatellites, in: Phylogeography and Population Genetics in Crustacea. CRC Press, Boca Raton, pp. 3–30.

Lucentini, L., Rebora, M., Puletti, M.E., Gigliarelli, L., Fontaneto, D., Gaino, E., Panara, F., 2011. Geographical and seasonal evidence of cryptic diversity in the Baetis rhodani complex (Ephemeroptera, Baetidae) revealed by means of DNA taxonomy. Hydrobiologia 673, 215–228. https://doi.org/10.1007/s10750-011-0778-1

Macher, J.N., Salis, R.K., Blakemore, K.S., Tollrian, R., Matthaei, C.D., Leese, F., 2016. Multiple-stressor effects on stream invertebrates: DNA barcoding reveals contrasting responses of cryptic mayfly species. Ecological Indicators 61, 159–169. https://doi.org/10.1016/j.ecolind.2015.08.024

Macher, J.-N., Vivancos, A., Piggott, J.J., Centeno, F.C., Matthaei, C.D., Leese, F., 2018. Comparison of environmental DNA and bulk-sample metabarcoding using highly degenerate cytochrome c oxidase I primers. Molecular Ecology Resources 18, 1456–1468. https://doi.org/10.1111/1755-0998.12940

Oksanen, J., Blanchet, G., Friendly, M., Kindt, R., Legendre, P., Dan, M., Minchin, P., O’Hara, B., Simpson, G., Salymos, P., Stevens, H., Eduard, S., Helene, W., 2019. vegan: Community Ecology Package. R package version 2.5-5.

Pauls, S.U., Lumbsch, H.T., Haase, P., 2006. Phylogeography of the montane caddisfly Drusus discolor: evidence for multiple refugia and periglacial survival. Molecular Ecology 15, 2153–2169. https://doi.org/10.1111/j.1365-294X.2006.02916.x

Pfrender, M.E., Hawkins, C.P., Bagley, M., Courtney, G.W., Creutzburg, B.R., Epler, J.H., Fend, S., Ferrington, L.C., Hartzell, P.L., Jackson, S., Larsen, D.P., Lévesque, C.A., Morse, J.C., Petersen, M.J., Ruiter, D., Schindel, D., Whiting, M., 2010. Assessing Macroinvertebrate Biodiversity in Freshwater Ecosystems: Advances and Challenges in DNA-based Approaches. The Quarterly Review of Biology 85, 319–340. https://doi.org/10.1086/655118

Phillips, J.D., Gillis, D.J., Hanner, R.H., 2019. Incomplete estimates of genetic diversity within species: Implications for DNA barcoding. Ecology and Evolution 9, 2996–3010. https://doi.org/10.1002/ece3.4757

R Development Core Team, 2008. R: A language and environment for statistical computing.

Ratnasingham, S., Hebert, P.D.N., 2007. bold: The Barcode of Life Data System (http://www.barcodinglife.org). Molecular Ecology Notes 7, 355–364. https://doi.org/10.1111/j.1471-8286.2007.01678.x

Reusch, T.B.H., Ehlers, A., Hämmerli, A., Worm, B., 2005. Ecosystem recovery after climatic extremes enhanced by genotypic diversity. Proc Natl Acad Sci U S A 102, 2826. https://doi.org/10.1073/pnas.0500008102

Reusch, T.B.H., Hughes, A.R., 2006. The emerging role of genetic diversity for ecosystem functioning: Estuarine macrophytes as models. Estuaries and Coasts 29, 159–164. https://doi.org/10.1007/BF02784707

Reynolds, L.K., McGlathery, K.J., Waycott, M., 2012. Genetic Diversity Enhances Restoration Success by Augmenting Ecosystem Services. PLOS ONE 7, e38397. https://doi.org/10.1371/journal.pone.0038397

Ribeiro, R., Lopes, I., 2013. Contaminant driven genetic erosion and associated hypotheses on alleles loss, reduced population growth rate and increased susceptibility to future stressors: an essay. Ecotoxicology 22, 889–899. https://doi.org/10.1007/s10646-013-1070-0

Smith, A.J., Bode, R.W., Kleppel, G.S., 2007. A nutrient biotic index (NBI) for use with benthic macroinvertebrate communities. Ecological Indicators 7, 371–386. https://doi.org/10.1016/j.ecolind.2006.03.001

Sturmbauer, C., Opadiya, G.B., Niederstätter, H., Riedmann, A., Dallinger, R., 1999. Mitochondrial DNA reveals cryptic oligochaete species differing in cadmium resistance. Molecular Biology and Evolution 16, 967–974. https://doi.org/10.1093/oxfordjournals.molbev.a026186

Theissinger, K., Röder, N., Allgeier, S., Beermann, A.J., Brühl, C.A., Friedrich, A., Michiels, S., Schwenk, K., 2019. Mosquito control actions affect chironomid diversity in temporary wetlands of the Upper Rhine Valley. Molecular Ecology 28, 4300–4316. https://doi.org/10.1111/mec.15214

Tsuji, S., Miya, M., Ushio, M., Sato, H., Minamoto, T., Yamanaka, H., 2019. Evaluating intraspecific genetic diversity using environmental DNA and denoising approach: A case study using tank water. Environmental DNA n/a. https://doi.org/10.1002/edn3.44

Turon, X., Antich, A., Palacín, C., Præbel, K., Wangensteen, O.S., 2019. From metabarcoding to metaphylogeography: separating the wheat from the chaff. Ecological Applications n/a. https://doi.org/10.1002/eap.2036

Usseglio-Polatera, P., Bournaud, M., Richoux, P., Tachet, H., 2000. Biological and ecological traits of benthic freshwater macroinvertebrates: relationships and definition of groups with similar traits. Freshwater Biology 43, 175–205. https://doi.org/10.1046/j.1365-2427.2000.00535.x

Van Dyck, H., 2012. Changing organisms in rapidly changing anthropogenic landscapes: the significance of the ‘Umwelt’-concept and functional habitat for animal conservation. Evolutionary Applications 5, 144–153. https://doi.org/10.1111/j.1752-4571.2011.00230.x

van Straalen, N.M., Timmermans, M.J.T.N., 2002. Genetic Variation in Toxicant-Stressed Populations: An Evaluation of the “Genetic Erosion” Hypothesis. Human and Ecological Risk Assessment: An International Journal 8, 983–1002. https://doi.org/10.1080/1080-700291905783

Vellend, M., 2005. Species Diversity and Genetic Diversity: Parallel Processes and Correlated Patterns. The American Naturalist 166, 199–215. https://doi.org/10.1086/431318

Vellend, M., Geber, M.A., 2005. Connections between species diversity and genetic diversity. Ecology Letters 8, 767–781. https://doi.org/10.1111/j.1461-0248.2005.00775.x

Vörösmarty, C.J., McIntyre, P.B., Gessner, M.O., Dudgeon, D., Prusevich, A., Green, P., Glidden, S., Bunn, S.E., Sullivan, C.A., Liermann, C.R., Davies, P.M., 2010. Global threats to human water security and river biodiversity. Nature 467, 555.

Wallace, J.B., Webster, J.R., 1996. The Role of Macroinvertebrates in Stream Ecosystem Function. Annu. Rev. Entomol. 41, 115–139. https://doi.org/10.1146/annurev.en.41.010196.000555

Weiss, M., Leese, F., 2016. Widely distributed and regionally isolated! Drivers of genetic structure in Gammarus fossarum in a human-impacted landscape. BMC Evolutionary Biology 16, 153. https://doi.org/10.1186/s12862-016-0723-z

Wickham, H., 2016. ggplot2: Elegant Graphics for Data Analysis. Springer-Verlag New York.

Williams, H.C., Ormerod, S.J., Bruford, M.W., 2006. Molecular systematics and phylogeography of the cryptic species complex Baetis rhodani (Ephemeroptera, Baetidae). Molecular Phylogenetics and Evolution 40, 370–382. https://doi.org/10.1016/j.ympev.2006.03.004

Witt, J.D., Hebert, P.D., 2000. Cryptic species diversity and evolution in the amphipod genus Hyalella within central glaciated North America: a molecular phylogenetic approach. Can. J. Fish. Aquat. Sci. 57, 687–698. https://doi.org/10.1139/f99-285

WWF, 2018. Living Planet Report 2018: Aiming higher. WWF, Gland, Switzerland.

Zizka, V.M.A., Elbrecht, V., Macher, J.-N., Leese, F., 2019. Assessing the influence of sample tagging and library preparation on DNA metabarcoding. Molecular Ecology Resources 19, 893–899. https://doi.org/10.1111/1755-0998.13018

