## Supplementary figures and images for "Can metabarcoding resolve intraspecific genetic diversity changes to environmental stressors? A test case using river macrozoobenthos"

### Supplemental Figure 1

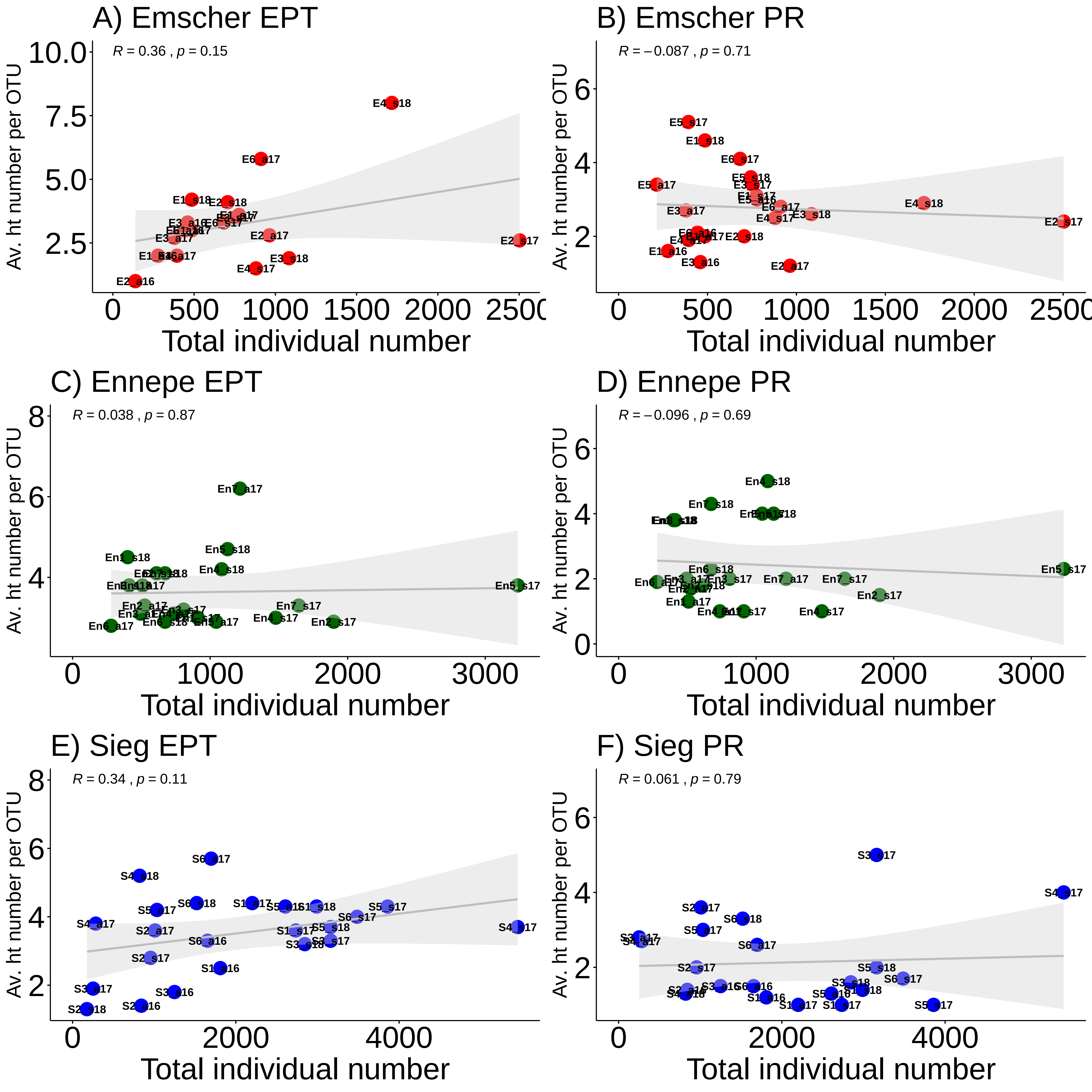

### Supplemental Figure 2

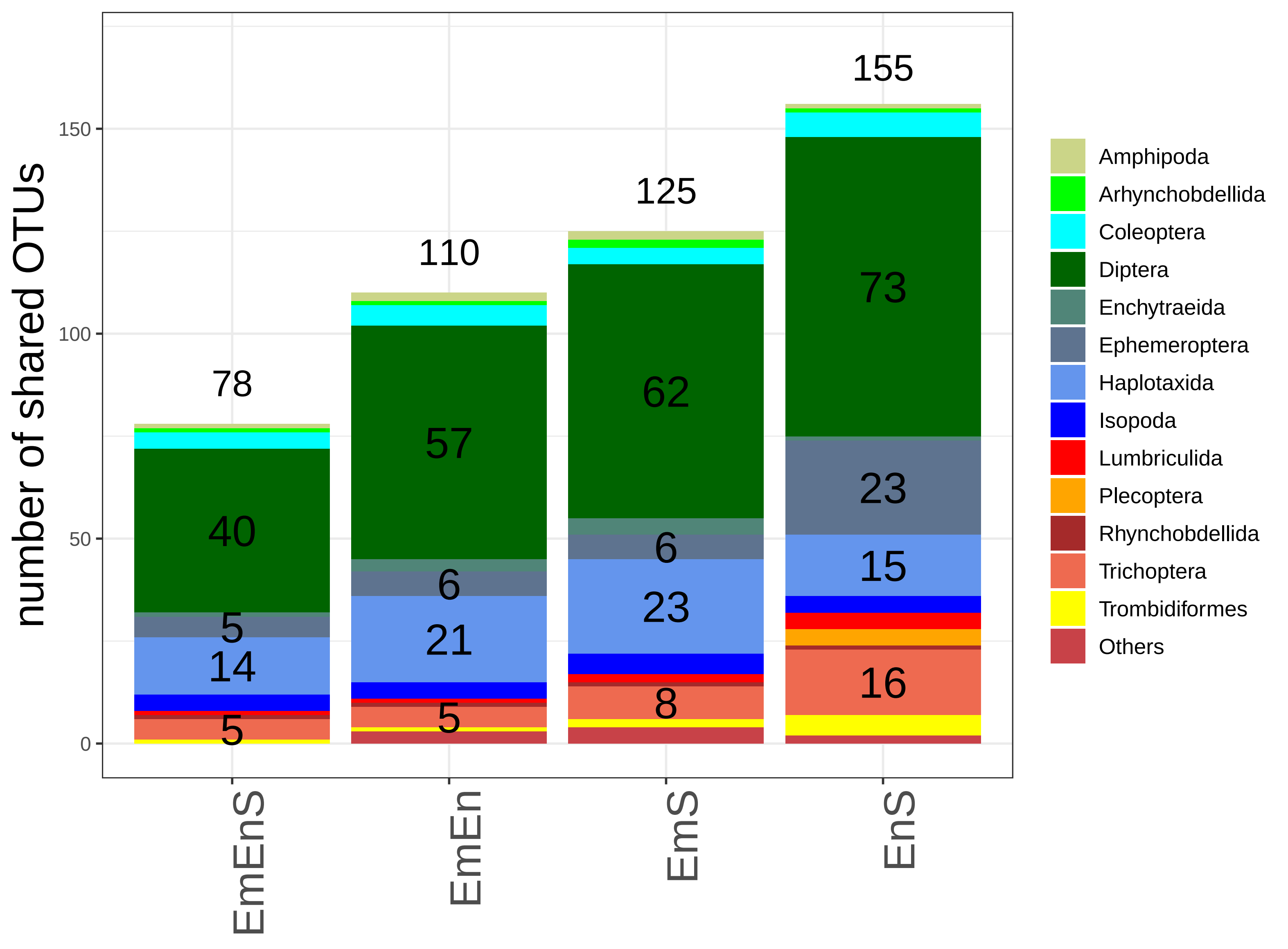

### Supplemental Figure 3

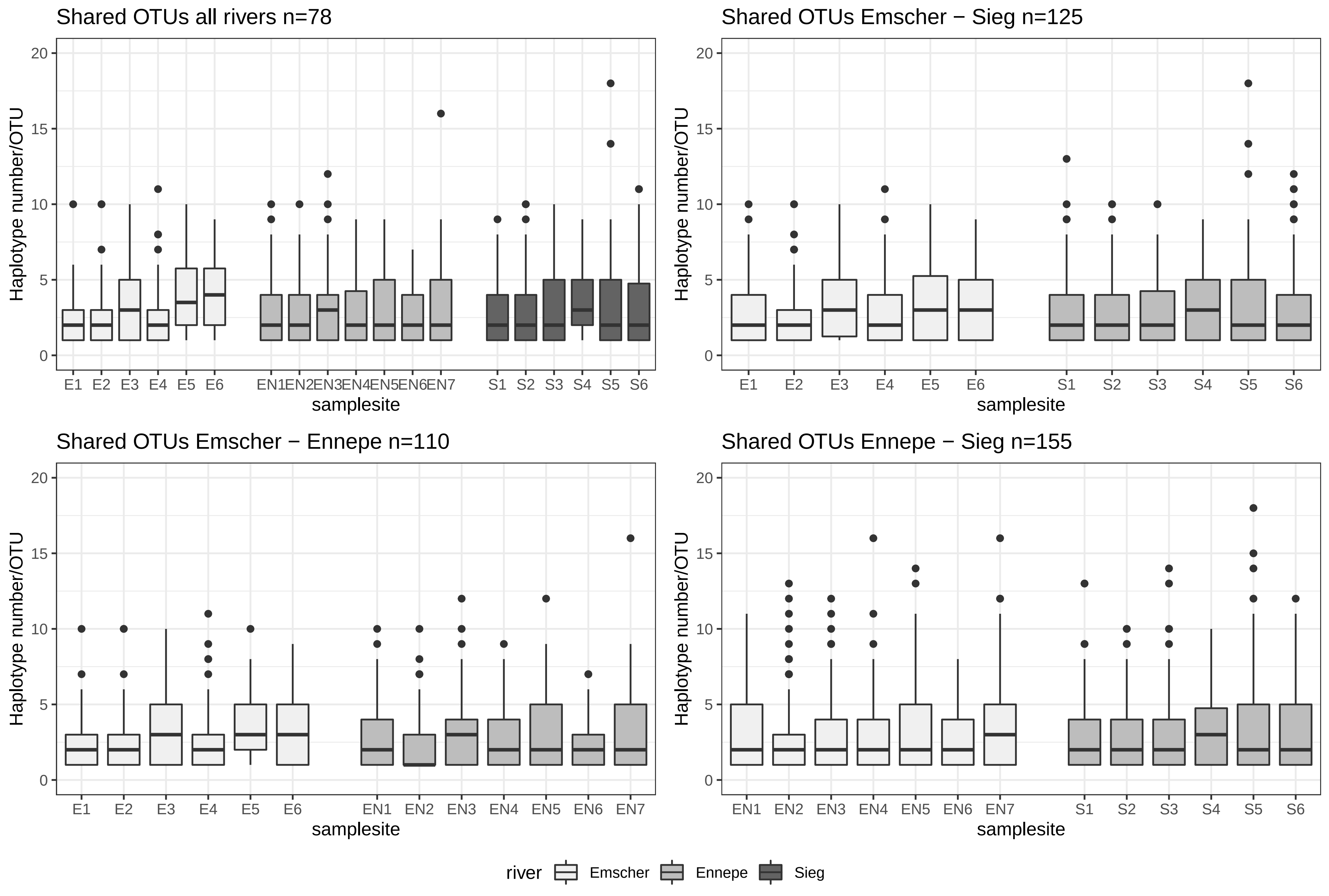

### Supplemental Figure 4

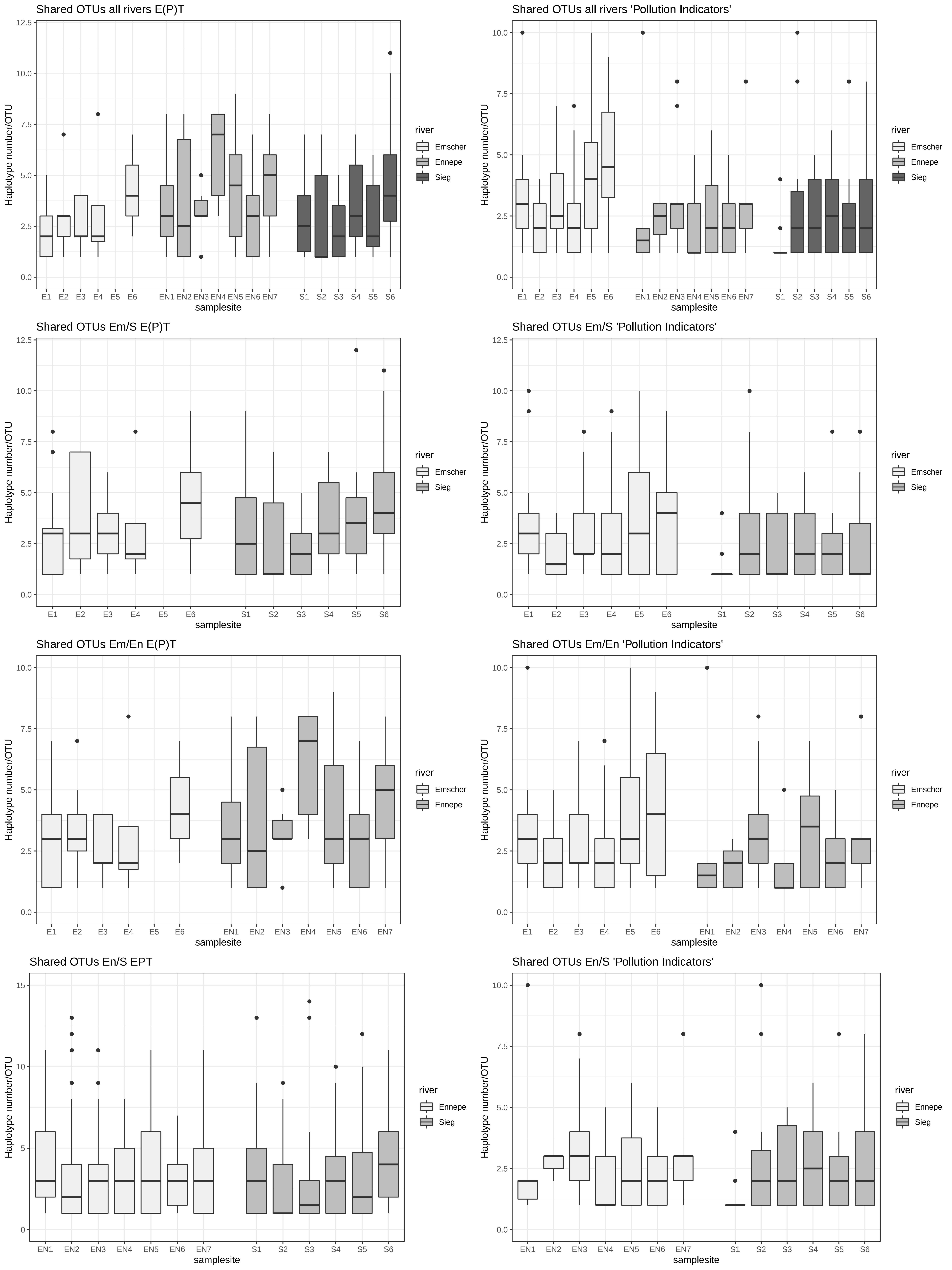

### Supplemental Figure 5

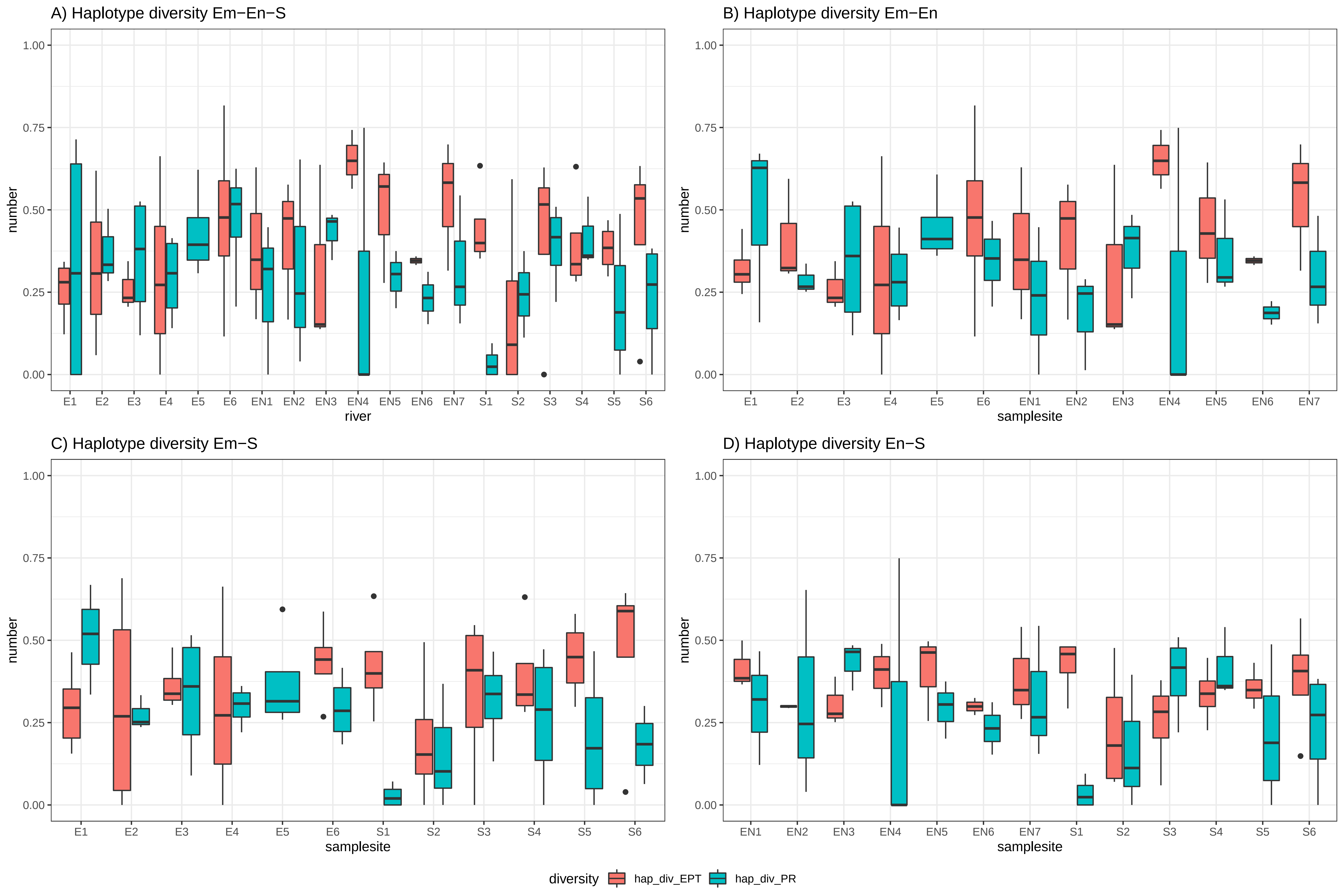

### Supplemental Figure 6

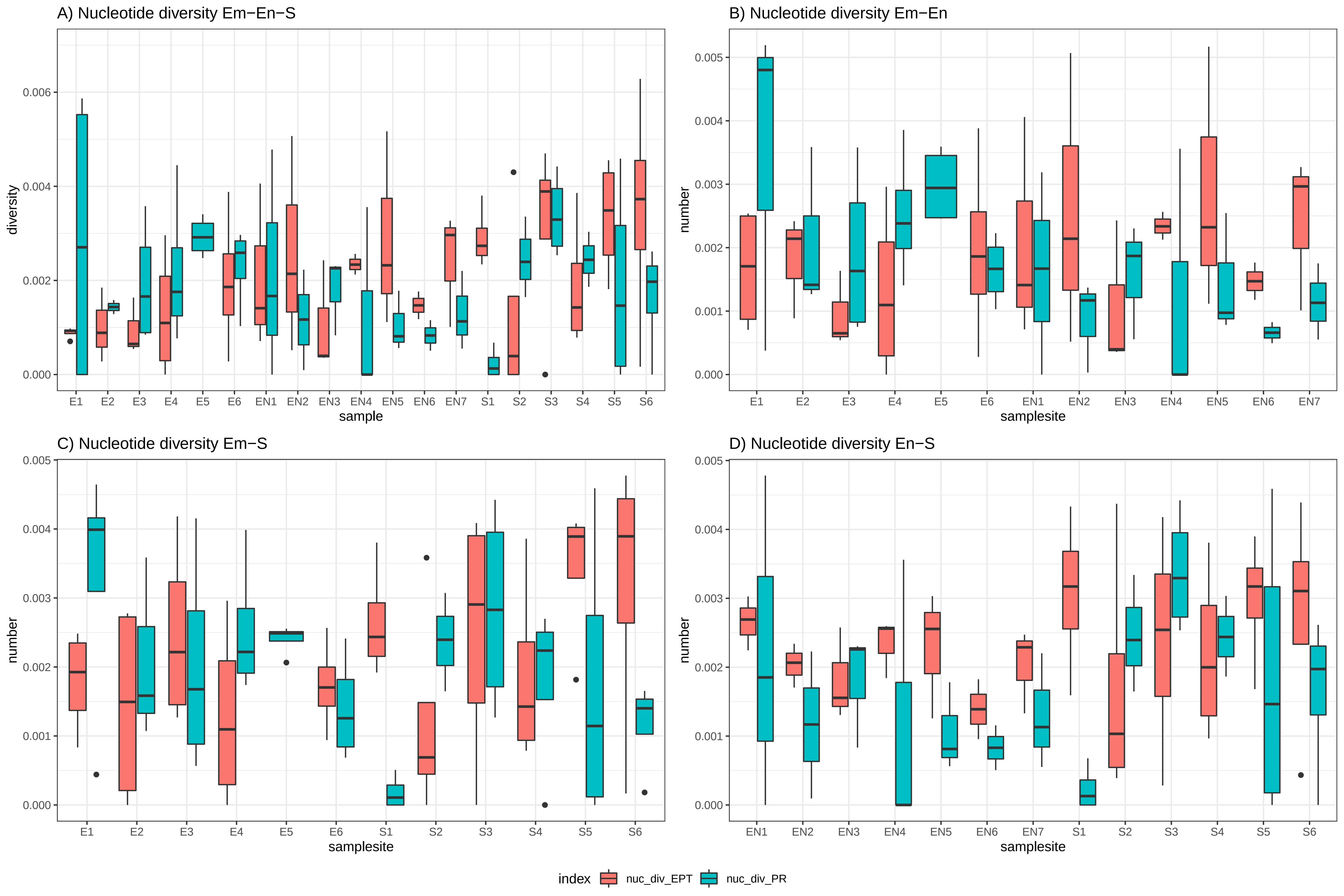

### Supplemental Figure 7

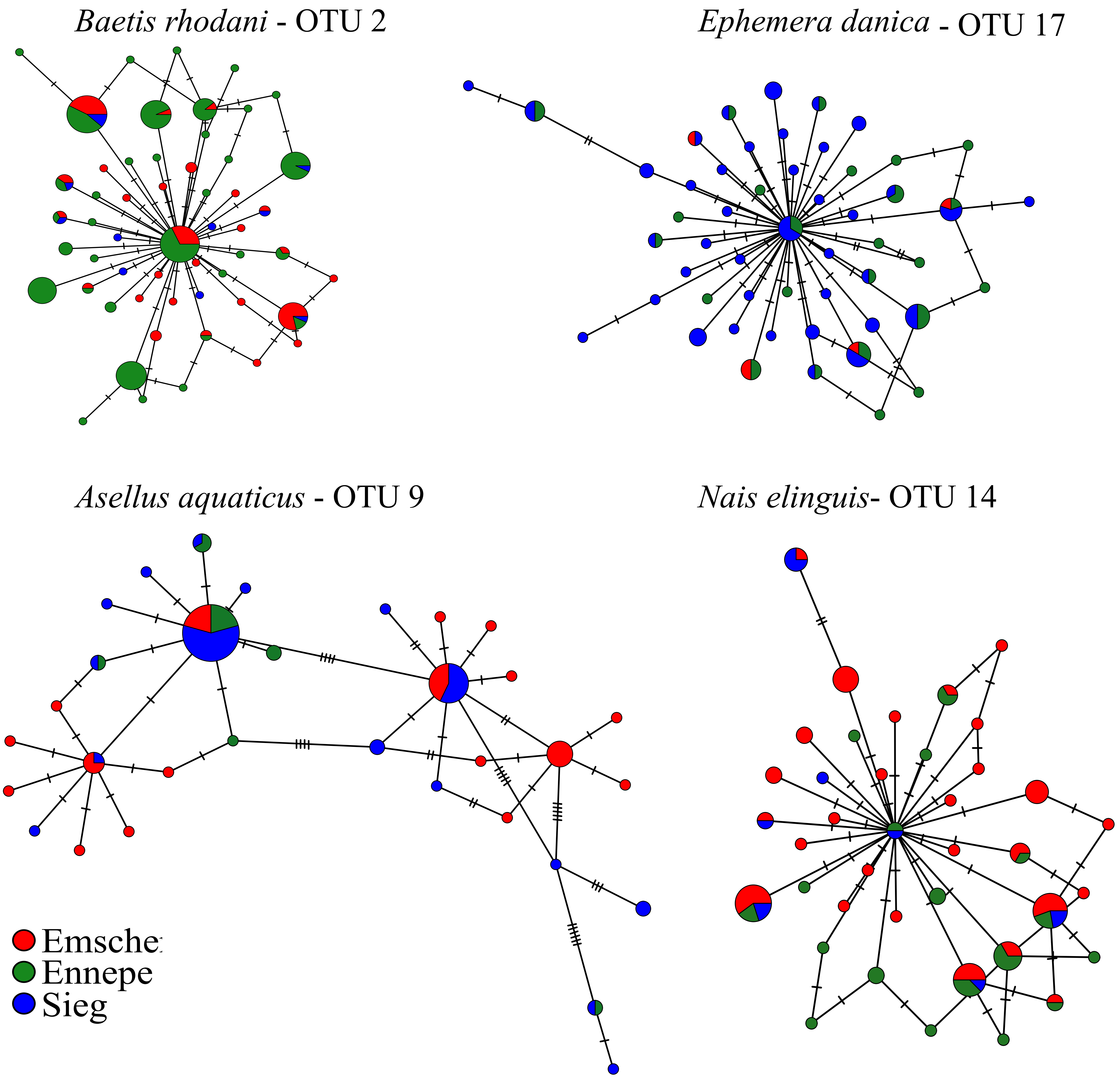
